# The road less travelled: Exploring the genomic characteristics and antimicrobial resistance potential of *Acinetobacter baumannii* from the indigenous Orang Asli community in Peninsular Malaysia

**DOI:** 10.1101/2025.06.12.659425

**Authors:** Soo-Sum Lean, Denise E. Morris, Rebecca Anderson, Ahmed Ghazi Alattraqchi, David W. Cleary, Stuart C. Clarke, Chew Chieng Yeo

**Author notes:** Corresponding authors: Stuart C. Clarke Chew Chieng Yeo.

## Abstract

*Acinetobacter baumannii* is widely recognized as a multidrug-resistant pathogen, but with most public genome datasets biased toward hospital-derived isolates. Little is thus known about *A. baumannii* isolates from healthy individuals from the community. This study analysed genome sequences from nine *A. baumannii* isolates obtained from the upper respiratory tract of healthy individuals of the indigenous Orang Asli community in their rural settlements located in the east coast of Peninsular Malaysia. Genomic analysis revealed all nine *A. baumannii* isolates to be genetically distinct and included three new sequence types (STs) under the Pasteur scheme and six novel STs under the Oxford scheme. Notably, one isolate, *A. baumannii* 19064, belonged to Global Clone 8 (GC8), a lineage associated with clinical infections. Core genome phylogeny indicated that the Orang Asli community isolates were interleaved with several non-GC clinical isolates from the tertiary hospital within the same state. This is suggestive of the potential of these community isolates to cause infections under conducive conditions. This is also attested by the identification of several virulence factors in their genomes. All nine isolates carried various intrinsic *bla*_OXA-51-like_ class D β-lactamase genes and class C *Acinetobacter*-derived cephalosporinase (ADC) *bla*_ADC_ genes but remained susceptible to meropenem. Two isolates, *A. baumannii* 19053 and 19062, were tetracycline resistant but minocycline susceptible and harboured the *tet(39)-tetR* gene pair within mobile p*dif* modules located on distinct Rep_3 family plasmids. Only one isolate, *A. baumannii* 19055, is plasmid-free; the rest mainly harboured cryptic Rep_3-type plasmids, often containing identifiable p*dif* modules. These findings highlight the clinical relevance of *A. baumannii* strains residing in healthy individuals, particularly in isolated communities that are seldom accessible to public health. Despite their remote location, the Orang Asli *A. baumannii* isolates possess virulence factors and antibiotic resistance genes similar to those found in hospital settings. This underscores the importance of genomic surveillance of commensal pathogens, and taking this road which is less travelled can help inform broader epidemiological insights and guide future public health strategies.

## 1 Introduction

Carbapenem-resistant *Acinetobacter baumannii* is recognised by the World Health Organization as the top critical priority pathogen, posing the highest threat to public health due to limited treatment options (World Health Organization, 2024). Not surprisingly, the majority of *A. baumannii* genomes currently reported and deposited in public databases, such as NCBI Genomes and the PubMLST genome collection, are mainly of hospital origin. These isolates are obtained from patients with hospital-acquired infections (HAIs), such as ventilator-associated pneumonia (VAP), meningitis, blood stream infections and urinary tract infections (Bian et al., 2021; Cui et al., 2023; Shelenkov et al., 2024). As a result, the public *A. baumannii* genome datasets are heavily skewed toward dominant hospital-associated clones, notably members of the notorious Global Clone 2 (GC2) lineage (Shelenkov et al., 2023). While keeping track of hospital-related *A. baumannii* is essential due to their formidable antimicrobial resistance (AMR) and clinical relevance, they only represent a subset of the species. These strains often differ significantly from those found in the normal flora of healthy populations (Muzahid et al., 2023). *A. baumannii* from healthy communities remain largely understudied, more so for indigenous communities that are isolated from urban populations.

The Orang Asli are indigenous people in Peninsular Malaysia comprising of several ethnic subgroups who retained their aboriginal language, customs and lifestyle (Mahmud et al., 2022). They are only a minor population in Malaysia (0.8% of the population in Peninsular Malaysia based on the year 2020 census) and often fall behind national socioeconomic, education and healthcare improvement plans. Despite government resettlement programmes, many Orang Asli communities remain in rural areas due to their lifestyle preferences (Syed Hussain et al., 2017; Mahmud et al., 2022). Some of these isolated communities are difficult to reach, limiting their access to modern medicines such as antibiotics and vaccines (Mohd Rosman et al., 2020; Chew et al., 2022). This restricted interaction with urban communities and healthcare has led to the formation of what we term here as “genomic capsules” – distinct microflora genomes unique to the Orang Asli community and their respective tribes. Consequently, opportunistic pathogens such as *A. baumannii* harboured by the Orang Asli may differ from strains commonly present in urban hospitals.

*A. baumannii* from hospitals have been well-studied over the past two decades, gaining their notoriety due to their multidrug resistance (MDR), extensive drug resistance (XDR) and pan-drug resistance (PDR) characteristics (Shi et al., 2024). The Malaysian Ministry of Health has published annual National Surveillance of Antibiotic Resistance (NSAR) Reports since 2003. Beginning in the 2010s, more than 50% of *A. baumannii* isolates have been reported to be resistant to carbapenems (i.e., imipenem and meropenem), which are the drugs of choice for treatment. However, the NSAR dataset is limited to participating hospitals which are mainly in the urban and suburban areas of Malaysia (Ministry of Health, 2023). While the World Health Organization’s (WHO) Tracking AMR Country Self-Assessment Survey (TrACSS) Country Report emphasizes the importance of addressing AMR at the community level to enhance infection prevention and control (IPC) efforts (WHO, 2022), there remains a significant knowledge gap regarding AMR profiles and genomic characteristics of bacterial pathogens within the indigenous communities in Malaysia. Notably, a recent study investigating *A. baumannii* isolates from human faecal samples in a community in Segamat, Malaysia, revealed phylogenomic clustering of four community-derived strains with two isolates from the town’s main tertiary hospital. This finding suggests the potential persistence and circulation of certain

*A. baumannii* strains across both community and healthcare settings (Muzahid et al., 2023). In this study, we aim to provide a genomic snapshot of *A. baumannii* that were isolated during an all-age, upper respiratory tract microbial carriage study undertaken among two rural Orang Asli communities in the state of Terengganu, located in the eastern coast of Peninsular Malaysia in 2017. A previous investigation of *Klebsiella pneumoniae* isolates from a broader indigenous cohort showed the predominance of ST23 which is commonly associated with clinical *K. pneumoniae* infections, and of concern, a proportion of these isolates harboured genes that categorised them as hypervirulent (Das et al., 2024). Here, we present the genomic analysis of *A. baumannii* isolates recovered from the upper respiratory tract of the Orang Asli and show the genetic diversity of this hitherto unexplored *A. baumannii* “genomic capsule”. Our findings offer insights into strains of novel STs, their patchwork of unique and shared mobile genetic elements, antimicrobial resistance and virulence genes.

## 2 Materials and Methods

### 2.1 Sampling and isolation of A. baumannii from Orang Asli communities

Swabs were taken from two Orang Asli villages, namely Kampung Sungai Pergam and Kampung Berua, in the state of Terengganu on the east coast of Peninsular Malaysia. Nasal swabs and nasopharyngeal swabs were taken from each participant, as described (Cleary et al., 2021). Conventional bacteriology of the samples were carried out using Columbia Blood Agar (CBA), CHOC agar (CBA with chocolated horse blood), CNA agar (CBA with colistin and naladixic acid), BACH (CBA with chocolated horse blood and Bacitracin) and GC agar (Lysed GC selective agar) (Cleary et al., 2021). Preliminary identification of presumptive *Acinetobacter* spp. isolates was done using the MALDI Biotyper (Bruker, UK) at the Portsmouth Microbiology Laboratories, UK.

### 2.2 Antibiotic susceptibility tests

*A. baumannii* was spread over Mueller-Hinton agar plates (MH; Oxoid, UK). Susceptibilities to the antibiotics meropenem and ciprofloxacin were determined by disk diffusion using the appropriate antibiotic disk (Oxoid, UK) whereas tetracycline and doxycycline susceptibilities were determined by placing Minimum inhibitory concentration (MIC) E-test strips (bioMérieux, France) onto the surface of the agar. All agar plates were incubated at 35°C ± 1°C for 18 h ± 2 h. Susceptibility was determined against the EUCAST Clinical Breakpoint guidelines (2024).

### 2.3 Genome Sequencing and Assemblies

Genomic DNA of the nine *A. baumannii* isolates were extracted using the QIAmp DNA Mini extraction kit (Qiagen, UK) per the manufacturer’s instructions. Concentration of genomic DNA was determined using Qubit 2.0 fluorometer (Thermo-Fisher, UK). Whole genome sequencing was performed on a MiSeq (Illumina, UK) short-read platform at a commercial sequencing provider (MicrobesNG, UK) using the 500 cycle v2 reagent kit to generate 2×150 bp paired-end reads. Raw reads obtained were then quality assessed and trimmed using fastp (available from https://github.com/OpenGene/fastp; (Chen, 2023)). Genome assemblies were carried out using Unicycler (available from https://github.com/rrwick/Unicycler; (Wick et al., 2017), followed by evaluation using Quast (available from https://github.com/ablab/quast).

### 2.4 Bioinformatics Analyses

Average nucleotide identity (ANI) of the assembled genomes to reference genomes were determined using fastANI (available from https://github.com/ParBLiSS/FastANI; (Jain et al., 2018). Annotation of the genomes was performed using Prokka (available from https://github.com/tseemann/prokka; (Seemann, 2014)). Conventional multilocus sequence typing (MLST) profiles of the assembled genomes were determined using mlst (available from https://github.com/tseemann/mlst) and matched to the PubMLST database (https://pubmlst.org/organisms/acinetobacter-baumannii). MLST profiles determined from the two available schemes, namely the Oxford and Pasteur schemes, were used to identify the corresponding Global Clones (GC). Serotyping based on *A. baumannii* surface polysaccharide loci, namely capsule K loci (KL) and lipo-oligosaccharide OC loci (OCL), were carried out using Kaptive v3.0.0b6 (available from https://github.com/klebgenomics/Kaptive; (Wyres et al., 2020; Cahill et al., 2022)).

Genotypic resistance profiles of the genomes were determined using AMRFinderPlus (available from https://github.com/ncbi/amr; (Feldgarden et al., 2021)) and ABRicate (available from https://github.com/tseemann/abricate), whereby databases from CARD (Alcock et al., 2023) and ResFinder (Zankari et al., 2012) were utilised to the latter approach. Virulome of the assembled genomes were determined using ABRicate, utilising database from Virulence Factor DataBase (VFDB) (Liu et al., 2022). Findings were then compared to the results obtained from VFAnalyzer (available from https://www.mgc.ac.cn/cgi-bin/VFs/v5/main.cgi). MGEs such as plasmids, insertion sequence (IS) elements and resistance island (RI) hotspots were also determined from the genomes. Plasmids were identified using PlasmidFinder (available from https://github.com/genomicepidemiology/plasmidfinder; (Carattoli et al., 2014)), whereas classification of the plasmid replication protein (Rep) was performed using an in-house built script, pREPonly (https://github.com/lean-SS/pREP-only) utilizing the AcinetobacterPlasmidTyping database (available from https://github.com/MehradHamidian/AcinetobacterPlasmidTyping; (Lam et al., 2023)). p*dif* sites in the plasmids found were identified using a combination of p*dif* finder (https://github.com/mjshao06/pdifFinder) (Shao et al., 2023) and manual search as outlined by Ambrose and Hall (2024). Toxin-antitoxin (TA) systems were determined using the TADB 3.0 database (https://bioinfo-mml.sjtu.edu.cn/TADB3/index.php) (Guan et al., 2024). IS elements were screened using ISEScan (available from https://github.com/xiezhq/ISEScan; (Xie and Tang, 2017)) and ISfinder-sequences database through the Prokka *–protein* option (available from https://github.com/thanhleviet/Isfinder-sequences; (Siguier et al., 2006)), whereas the *comM* resistance island (RI) hotspot (Hamidian and Hall, 2017) was determined through local BLAST.

### 2.5 Pangenome and Phylogenetic Analysis

The pangenomes of the nine Orang Asli *A. baumannii* isolates were compared to other community *A. baumannii* genomes using the Anvi’o platform (available from https://github.com/merenlab/anvio; (Delmont and Eren, 2018; Eren et al., 2021)). Through the use of the *anvi-pan-genome* programme, the pangenomes were determined and then visualised through *anvi-display-pan*, which links to the Anvi’o server. Core genome phylogenetic analysis of *A. baumannii* genomes were performed using Roary (available from https://sanger-pathogens.github.io/Roary/; (Page et al., 2015)) and VeryFastTree with the GTR+CAT model, which combines the General Time Reversible (GTR) nucleotide substitution model with a Constant Rate Across Sites (CAT) approximation (Piñeiro et al., 2020; Piñeiro and Pichel, 2024), and visualised with iTOL v7 (Letunic and Bork, 2024).

## 3.0 Results and Discussions

### 3.1 Preliminary Genomic Analysis of A. baumannii from the Orang Asli

A total of thirteen presumptive *Acinetobacter* spp. isolates were obtained from the Orang Asli carriage studies at Kampung Sungai Pergam (*n* = 3) and Kampung Berua (*n* = 10). Whole genome sequencing was performed on all thirteen isolates, out of which nine were shown to be *A. baumannii* (Table 1). The remaining four isolates were determined to be *A. nosocomialis*.

**Table 1:** General information of the source and the genome characteristics of the *A. baumannii* (*n* = 9) isolates from the upper respiratory tract of the Orang Asli in this study.

| *Acinetobacter* isolate | 19053 | 19055 | 19056 | 19058 | 19060 | 19061 | 19062 | 19063 | 19064 |
| --- | --- | --- | --- | --- | --- | --- | --- | --- | --- |
| Location/source<sup>§</sup> | KSP/N | KSP/NP | KB/N | KB/N | KB/NP | KB/N | KB/NP | KB/N | KB/N |
| Species identification | *Acinetobacter baumannii* | *Acinetobacter baumannii* | *Acinetobacter baumannii* | *Acinetobacter baumannii* | *Acinetobacter baumannii* | *Acinetobacter baumannii* | *Acinetobacter baumannii* | *Acinetobacter baumannii* | *Acinetobacter baumannii* |
| Accession no. | JBNPBH<br>000000000 | JBNPBI<br>000000000 | JBNPBJ<br>000000000 | JBNPBK<br>000000000 | JBNPBL<br>000000000 | JBNPBM<br>000000000 | JBNPBN<br>000000000 | JBNPBO<br>000000000 | JBNPBP<br>000000000 |
| Closest neighbour (fastANI) | *Acinetobacter baumannii*<br>SAMEA<br>104305267<br>(98.1849%) | *Acinetobacter baumannii*<br>5577<br>STDY7716391<br>(99.8833%) | *Acinetobacter baumannii*<br>5577<br>STDY7716201<br>(97.9228%) | *Acinetobacter baumannii*<br>5577<br>STDY7715392<br>(98.7137%) | *Acinetobacter baumannii*<br>SAMEA<br>104305309<br>(98.8467%) | *Acinetobacter baumannii*<br>SAMEA<br>104305313<br>(98.0632%) | *Acinetobacter baumannii*<br>SAMN<br>03174920<br>(98.6532%) | *Acinetobacter baumannii*<br>SAMN<br>03174917<br>(97.9238%) | *Acinetobacter baumannii*<br>SAMEA<br>104305269<br>(99.8239%) |
| Total genome size (bp) | 3,821,374 | 3,697,509 | 3,748,932 | 3,904,855 | 3,767,717 | 3,865,738 | 3,844,544 | 3,828,269 | 3,664,686 |
| No. of contigs | 62 | 166 | 32 | 44 | 126 | 35 | 149 | 47 | 47 |
| No. of plasmids | 2 | 0 | 1 | 1 | 1 | 1 | 2 | 1 | 2 |
| GC content (%) | 38.91 | 39.04 | 38.97 | 38.87 | 38.95 | 38.75 | 38.99 | 38.81 | 38.78 |
| N<sub>50</sub> | 478,783 | 67,766 | 458,370 | 254,121 | 77,219 | 477,098 | 73,853 | 369,580 | 195,600 |
| N<sub>90</sub> | 68,035 | 15,343 | 166,877 | 88,568 | 17,967 | 165,195 | 17,420 | 116,120 | 67,830 |
| L<sub>50</sub> | 3 | 17 | 3 | 6 | 16 | 2 | 14 | 3 | 7 |
| L<sub>90</sub> | 11 | 61 | 8 | 18 | 49 | 8 | 53 | 10 | 20 |
| No. of CDS | 3,579 | 3,428 | 3,497 | 3,661 | 3,582 | 3,573 | 3,674 | 3,576 | 3,490 |
| MLST<sub>Oxf</sub> | ST3542* | ST3543* | ST3415 | ST3541* | ST871 | ST3544* | ST3545* | ST3540* | ST585 |
| MLST<sub>Pas</sub> | ST2832* | ST2114 | ST2522 | ST2834* | ST2700 | ST470 | ST2635 | ST2833* | ST10 |
| K loci | KL121 | KL96 | KL170<sup>#</sup> | KL202<sup>#</sup> | KL112 | KL172<sup>#</sup> | KL81 | KL183<sup>#</sup> | KL108 |
| OC loci | OCL2 | OCL13 | OCL2 | OCL4 | OCL6 | OCL4 | OCL6 | OCL2 | OCL2 |

<sup>§</sup>Abbreviations used for location/source: KSP, Kampung Sungai Pergam; KB, Kampung Berua; N, nasal swab; NP, nasopharyngeal swab.
Notes: MLST<sub>Oxf</sub> and MLST<sub>Pas</sub> represent the Oxford and Pasteur schemes available in PubMLST, respectively. ST represents sequence type and those marked with an asterisk (\*) are new STs that were identified in this study; KL represents K loci with those labelled with <sup>#</sup> representing K loci that has not been reported; OCL represents lipo-oligosaccharide OC loci.

Genome assemblies of the nine *A. baumannii* Orang Asli isolates showed total genome sizes that ranged from ∼3.7 Mbp to 3.9 Mbp (**Table 1**). Average nucleotide identity (ANI) of the genomes revealed >97% nucleotide identities to *A. baumannii* genomes including those from Thailand (SAMEA104305267, SAMEA104305309, SAMEA104305313 and SAMEA104305269), Vietnam (5577STDY7716391, 5577STDY7716201 and 5577STDY7716392) and Malaysia (SAMN03174920 and SAMN03174917), as listed in **Table 1**. *In silico* epidemiological typing of the assembled genomes, which includes traditional MLST and surface polysaccharide loci typing, showed that each genome is distinct with its own sequence type (ST), KL and OCL types. MLST profiles based on the Oxford scheme revealed six new STs, with only *A. baumannii* 19056, 19060 and 19064 that were identified as preexisting ST3415_Oxf_, ST871_Oxf_ and ST585_Oxf_, respectively (**Table 1**). The new Oxford STs were submitted and assigned by the PubMLST curators as ST3540_Oxf_, ST3541_Oxf_, ST3542_Oxf_, ST3543_Oxf_, ST3543_Oxf_, ST3544_Oxf_ and ST3545_Oxf_ (**Table 1**). Additionally, the Pasteur MLST scheme identified three novel STs from the nine *A. baumannii* genomes, and these were assigned as ST2832_Pas_, ST2833_Pas_, and ST2844_Pas_. The six remaining genomes each belonged to different preexisting Pasteur STs (**Table 1**).

*A. baumannii* surface polysaccharide loci typing based on the K loci (KL), which were responsible for the production of acinetamic acid (Lam et al., 2022), showed that each of the nine Orang Asli *A. baumannii* genomes belonged to distinct KL types (**Table 1**). Notably, four novel capsule types (i.e., KL170, KL202, KL172 and KL183) were detected, indicating previously unreported characteristics and underscoring the uniqueness of these Orang Asli *A. baumannii* isolates. In contrast, analysis of the outer core lipo-oligosaccharide loci (OCL) showed OCL2 (*n* = 4) to be the most common type, followed by OCL4 (*n* = 2) and OCL6 (*n* = 2) (**Table 1**). Nevertheless, *A. baumannii* 19055 was identified as the lesser-studied OCL13 type, which was originally described in *A. baumannii* strains associated with community-acquired pneumonia in the Northern Territories, Australia (Meumann et al., 2019).

### 3.2 Genotypic AMR observation revealed non-mainstream β-lactamases

The presence of genes encoding β-lactamases (*bla*) has become the hallmark of *A. baumannii*, not only as key determinants of carbapenem resistance (Shi et al., 2024), but also through the intrinsic *bla*_OXA-51-like_ genes, which have been adopted as one of the typing methods for *A. baumannii* over the past two decades (Shelenkov et al., 2023). The recent categorization of global clones (GCs) also incorporated various *bla* genes as characteristic traits observed in certain GCs, for example, members of the GC8 lineage commonly harbour both *bla*_OXA-23_ and *bla*_OXA-68_ (Shelenkov et al., 2023). Although *bla* genes such as *bla*_OXA-23_, *bla*_OXA-51_ and *bla*_OXA-66_, are widespread among *A. baumannii* globally (Li et al., 2023; Shelenkov et al., 2023; Shi et al., 2024), a different spectrum of *bla* genes were found from the Orang Asli *A. baumannii* in this study (**Table 2**).

**Table 2:** Distribution of antimicrobial resistance determinants across the Malaysian Orang Asli *A. baumannii* genomes (*n* = 9) and their corresponding phenotypic resistance profiles.

| Antimicrobial resistance genes/phenotype | *Acinetobacter baumannii* isolate |  |  |  |  |  |  |  |  |
| --- | --- | --- | --- | --- | --- | --- | --- | --- | --- |
|  | 19053 | 19055 | 19056 | 19058 | 19060 | 19061 | 19062 | 19063 | 19064 |
| **Beta-lactams:** |  |  |  |  |  |  |  |  |  |
| • Resistance gene(s) | *bla*<sub>ADC-25</sub>,<br>*bla*<sub>OXA-377</sub> | *bla*<sub>ADC-279</sub>,<br>*bla*<sub>OXA-120</sub> | *bla*<sub>ADC-169</sub> variant,<br>*bla*<sub>OXA-555</sub> | *bla*<sub>ADC-312</sub> variant,<br>*bla*<sub>OXA-51</sub> | *bla*<sub>ADC-238</sub>,<br>*bla*<sub>OXA-510</sub> | *bla*<sub>ADC-25</sub>,<br>*bla*<sub>OXA-51</sub> | *bla*<sub>ADC-99</sub>,<br>*bla*<sub>OXA-98</sub> | *bla*<sub>ADC-158</sub> variant,<br>*bla*<sub>OXA-424</sub> | *bla*<sub>ADC-76</sub>,<br>*bla*<sub>OXA-68</sub> |
| • Meropenem* | 22.3 (S) | 26.6 (S) | 24.2 (S) | 29.1 (S) | 28.6 (S) | 25.7 (S) | 21.0 (S) | 27.0 (S) | 24.6 (S) |
| **Aminoglycoside resistance gene(s)** | *ant(3'')-IIa* | *ant(3'')-IIa* | - | - | *ant(3'')-IIa* | - | *ant(3'')-IIa* | - | *ant(3'')-IIa* |
| **Tetracyclines:** |  |  |  |  |  |  |  |  |  |
| • Resistance gene(s) | *tet(39)* | - | - | - | - | - | *tet(39)* | - | - |
| • Tetracycline<sup>#</sup> | 64.0 (R) | 12.0 (S) | 4.0 (S) | 6.0 (S) | 6.0 (S) | 4.0 (S) | 128.0 (R) | 4.0 (S) | 6.0 (S) |
| • Doxycycline<sup>#</sup> | 4.0 (S) | 4.0 (S) | 4.0 (S) | 4.0 (S) | 4.0 (S) | 4.0 (S) | 4.0 (S) | 4.0 (S) | 4.0 (S) |
| **Sulphonamide resistance genes(s)** | - | - | - | - | - | - | *sul2* | - | - |
| **Fluroquinolones:** |  |  |  |  |  |  |  |  |  |
| • Mutation(s) in QRDRs of *gyrA* and *parC* | - | - | - | - | - | - | - | - | - |
| • Ciprofloxacin* | 23.5 (S) | 25.2 (S) | 24.7 (S) | 28.3 (S) | 28.0 (S) | 28.0 (S) | 21.7 (S) | 26.8 (S) | 24.8 (S) |
| **Efflux pumps** | *amvA*<br>*abeS*, *abeM*,<br>*adeA*, *adeF*,<br>*adeG*, *adeH*,<br>*adeI*, *adeJ*,<br>*adeK*, *adeL*,<br>*adeN* | *amvA*<br>*abeS*, *abeM*,<br>*adeF*, *adeG*,<br>*adeH*, *adeI*, *adeJ*,<br>*adeK*, *adeL*,<br>*adeN*, *adeS*, *adeR* | *amvA*<br>*abeS*, *abeM*,<br>*adeA*, *adeB*,<br>*adeF*, *adeG*,<br>*adeH*, *adeI*, *adeJ*,<br>*adeK*, *adeL*,<br>*adeN*, *adeS*, *adeR* | *amvA*<br>*abeS*, *abeM*,<br>*adeA*, *adeB*,<br>*adeF*, *adeG*,<br>*adeH*, *adeI*, *adeJ*,<br>*adeK*, *adeL*,<br>*adeN*, *adeS*, *adeR* | *amvA*<br>*abeS*, *abeM*,<br>*adeA*, *adeB*,<br>*adeF*, *adeG*,<br>*adeH*, *adeI*, *adeJ*,<br>*adeK*, *adeL*,<br>*adeN*, *adeS*, *adeR* | *amvA*<br>*abeS*, *abeM*,<br>*adeA*, *adeB*,<br>*adeF*, *adeG*,<br>*adeH*, *adeI*, *adeJ*,<br>*adeK*, *adeL*,<br>*adeN*, *adeS*, *adeR* | *amvA*<br>*abeS*, *abeM*,<br>*adeF*, *adeG*,<br>*adeH*, *adeI*,<br>*adeJ*, *adeK*,<br>*adeL*, *adeN* | *amvA*<br>*abeS*, *abeM*,<br>*adeA*, *adeB*,<br>*adeC*, *adeF*,<br>*adeG*, *adeH*,<br>*adeI*, *adeJ*, *adeK*,<br>*adeL*, *adeN* | *amvA*<br>*abeS*, *abeM*,<br>*adeB*, *adeC*,<br>*adeF*, *adeG*,<br>*adeH*, *adeI*,<br>*adeJ*, *adeK*,<br>*adeL*, *adeN* |

\*values indicate zone of inhibition (in mm) of respective antibiotic discs with interpretations of resistance (R) or susceptibility (S) following EUCAST (2024) breakpoints.
<sup>#</sup>values indicate minimum inhibitory concentrations (in µg/ml) measured using MIC E-test strips with interpretations of resistance (R) or susceptibility (S) following EUCAST (2024) breakpoints.

The *bla*_OXA-51_ family (or *bla*_OXA-51-like_) is comprised of the well-characterized *bla*_OXA-51_ gene and its numerous variants, such as *bla*_OXA-64_, *bla*_OXA-66_ and others (Li et al., 2023). In this study, several variants were detected in the Orang Asli *A. baumannii* genomes including *bla*_OXA-68_, *bla*_OXA-98_, *bla*_OXA-120_, *bla*_OXA-377_, *bla*_OXA-424_, *bla*_OXA-510_, and *bla*_OXA-555_. Only two genomes (i.e., *A. baumannii* 19058 and 19061) carried *bla*_OXA-51_ itself while the remaining seven harboured distinct variants of the *bla*_OXA-51_ family (**Table 2** and **Figure 3**). The spectrum of genes of the *bla*_OXA-51_ family aligns with the findings of (Muzahid et al., 2023) who reported that community-derived *A. baumannii* isolates from Segamat, Peninsular Malaysia also carried a wide range of *bla*_OXA-51_ variants. In the Segamat *A. baumannii* isolates, *bla*_OXA-120_ was the most prevalent variant, followed by *bla*_OXA-441_, *bla*_OXA-510_, *bla*_OXA-69_, *bla*_OXA-98_ and *bla*_OXA-412_ (Muzahid et al., 2023). Interestingly, this study found none of these variants among the *A. baumannii* isolates from the Orang Asli community.

The presence of the intrinsic *bla*_OXA-51_ family of genes is, however, not a marker for carbapenem resistance in *A. baumannii* due to the low affinities of the OXA-51 family of β-lactamases to meropenem and imipenem as well as their very low expression levels. In certain cases, the presence of IS*Aba1* directly upstream of the *bla*_OXA-51-like_ gene provides a strong promoter which increases its expression level, leading to carbapenem resistance but this also depends on the *bla*_OXA-51_ variant that is being overexpressed (Nigro and Hall, 2018). All nine Orang Asli *A. baumannii* isolates were phenotypically carbapenem susceptible, and none of their genomes contained IS*Aba1* (or related elements) upstream of the *bla*_OXA-51-like_ genes. None of the nine Orang Asli *A. baumannii* isolates also harboured acquired *bla*_OXA_-encoded carbapenemase genes such as *bla*_OXA-23_, *bla*_OXA-24_, and *bla*_OXA-58_, which have been directly implicated in carbapenem resistance, particularly in clinical *A. baumannii* isolates (Hamidian and Nigro, 2019). This was similarly reported by (Muzahid et al., 2023) where the acquired carbapenemase gene *bla*_OXA-23_ were only identified in their Segamat hospital isolates which are carbapenem resistant. Likewise, our recent study of a 10-year collection of *A. baumannii* from the main tertiary hospital in Terengganu also showed the predominance of the *bla*_OXA-23_ gene among the carbapenem-resistant isolates (Din et al., 2025).

Another class of intrinsic β-lactamase found in *A. baumannii* genomes is the AmpC cephalosporinase variants which are designated *Acinetobacter*-derived cephalosporinases (ADCs). Overproduction of ADCs resulting from insertion of IS*Aba1* or similar IS elements upstream of the *bla*_ADC_ gene has been shown to be responsible for the development of resistance towards extended-spectrum cephalosporins and in some cases, carbapenems (Tian et al., 2011; Bhattacharya et al., 2014; Shi et al., 2024). Seven different variants of ADCs were detected from the genomes of the nine Orang Asli *A. baumannii* isolates (**Table 2**) and in all cases, IS*Aba1* or similar elements were absent upstream of the encoding gene, suggesting that these genes were either not expressed or were expressed at low levels in their hosts. ADC-25 were identified in two of the Orang Asli *A. baumannii* isolates (i.e., 19053 and 19061; **Table 2**) and this variant was found to be the 7^th^ most prevalent ADC variant among *A. baumannii* isolates globally (Mack et al., 2025). Four of the eight ADC variants identified here (i.e., ADC-99 in *A. baumannii* 19062, ADC-238 in 19060, ADC-279 in 19055, and ADC-312 in 19058) were listed by (Mack et al., 2025) as variants that were rarely found in *A. baumannii*. In comparison, the *A. baumannii* isolates from the Segamat community also presented a different spectrum of *bla*_ADC_ genes (Muzahid et al., 2023). The Segamat community isolates mainly harboured *bla*_ADC-154_ and *bla*_ADC-156_ (Muzahid et al., 2023) and both variants were absent in our Orang Asli isolates. However, *bla*_ADC-238_ which was found in *A. baumannii* C-65 from the Segamat community, was also found in *A. baumannii* 19060 form our Orang Asli collection. The ADC-238 variant was listed as a less-frequently encountered variant (Mack et al., 2025) and its singular presence in both these community-based studies attested to this finding. Intriguingly, hospital isolates from Segamat (Muzahid et al., 2023) and Terengganu (Din et al., 2025) showed a uniform pattern of *bla*_ADC-73_ being the most prevalent, agreeing with the analysis presented by (Mack et al., 2025) which revealed *bla*_ADC-73_ as the most prevalent ADC variant in *A. baumannii* isolates globally, with the exception of isolates from North America.

Hence, in terms of the class C (ADC) and class D (OXA) β-lactamases, the intrinsic variants harboured by the *A. baumannii* community isolates showed diversity with little in common although both studies (i.e., this study and (Muzahid et al., 2023)) were in Peninsular Malaysia. Even between communities, the AMR profiles varies (Meumann et al., 2019; Muzahid et al., 2023) and thus, there are more to be learned about *A. baumannii* from the Orang Asli community, which is further elaborated in the following sections.

### 3.3 Other resistance genes in the Orang Asli community A. baumannii isolates

Two out of the nine Orang Asli *A. baumannii* isolates (i.e., 19053 and 19062) were found to harbour the *tet(39)* tetracycline resistance gene (Table 2), which encodes a tetracycline efflux pump of the major facilitator superfamily (MFS) (Agersø and Guardabassi, 2005). This differs from the *A. baumannii* community isolates from Segamat in which no tetracycline resistance genes were detected. Both *A. baumannii* 19053 and 19062 were phenotypically tetracycline resistant (with MIC values of 64 µg/mL and 128 µg/mL, respectively) but minocycline susceptible (both with MIC values of 4 µg/mL). In contrast, Malaysian *A. baumannii* hospital isolates predominantly carried the *tet(B)* gene (*n* = 60/126 from HSNZ (Din et al., 2025); *n* = 12/15 from Segamat Hospital (Muzahid et al., 2023)), with only one isolate from HSNZ harbouring *tet(A)* and 10/126 carrying *tet(39)* (Din et al., 2025). All *A. baumannii* hospital isolates harbouring the *tet(39)* and *tet(A)* genes were tetracycline resistant and minocycline susceptible whereas those that harboured the *tet(B)* gene were mostly resistant to tetracycline but showed intermediate susceptibility to minocycline (Din et al., 2025). Meumann et al. (2019) reported two *A. baumannii* isolates from community-onset pneumonia in Australia which harboured both the *tet(B)* and *tet(39)* genes but phenotypic susceptibility testing for tetracyclines was not performed in their study. The two tetracycline-resistant Orang Asli *A. baumannii* isolates, 19053 and 19062, harboured the *tet(39)* gene on plasmids, which will be elaborated in a later section.

Resistance to aminoglycosides in *A. baumannii* is mainly mediated by the possession of genes encoding aminoglycoside acetyltransferase (*aac*), nucleotidyltransferase (*ant*) and/or phosphotransferase (*aph*) (Shi et al., 2024). Five out of the nine *A. baumannii* Orang Asli isolates carried the *ant(3”)-IIa* gene (Table 2) whereas Muzahid et al. (2023) reported the presence of this gene in all twelve of their *A. baumannii* strains that were isolates from the community in the town of Segamat. Nevertheless, only two of the twelve Segamat community isolates showed resistance to amikacin and all were gentamicin susceptible, suggesting that some of these aminoglycoside resistance genes were either not expressed or expressed at a very low level. Phenotypic resistance to aminoglycosides was, however, not tested for the Orang Asli *A. baumannii* isolates in this study.

Sulphonamide resistance in *A. baumannii* is usually mediated by *sul1* and/or *sul2* genes (Köld, 2001), with *sul2* predominantly reported from Southeast Asian countries and the Asia-Pacific region (Bian et al., 2021; Brito et al., 2022; Din et al., 2025). Only one Orang Asli isolate, *A. baumannii* 19062, was found to harbour the *sul2* gene (**Table 2**), which was absent in the Segamat community isolates (Muzahid et al., 2023). In contrast, nearly 50% (61/126) of the *A. baumannii* hospital isolates from HSNZ, Terengganu, harboured the *sul2* gene (Din et al., 2025) whereas in Hospital Segamat, the gene was identified in 2/15 of the *A. baumannii* isolates (Muzahid et al., 2023). The significance of the carriage of the *sul2* gene in the solitary Orang Asli *A. baumannii* isolate is currently unknown, but *sul2* is known to be present on mobile elements such as plasmids and transposons (Jeon et al., 2023). Plasmid analysis appeared to rule out the carriage of *sul2* in either of the two plasmids found in *A. baumannii* 19062 (see subsequent section 3.7) but this does not rule out its location on other mobile elements such as transposons or genomic islands in the chromosome.

### 3.4 The virulome of A. baumannii from the Orang Asli community

The isolation of *A. baumannii* from diverse sources, including clinical settings, soil, and wastewater, highlights its ability to persist across various environmental niches, facilitating its widespread dissemination, colonisation, and pathogenicity (Harding et al., 2018). This persistence, evidenced by stress resistance and biofilm formation, is likely supported by the acquisition and/or inheritance of multiple virulence factors (VFs). In this study, the majority of identified VFs are associated with adhesion, biofilm formation, and quorum sensing regulation (**Figure 1**). These findings are in agreement with Muzahid et al. (2023), who reported similar virulome profiles in *A. baumannii* isolates from the Segamat community, including genes related to adhesion (e.g., *ompA*, *fliP*, *pilA*, *pilE*), biofilm formation (e.g., *adeFGH*, *bap*, *csuA/B*, *csuABCDE*, *pgaABCD*), and quorum sensing (e.g., *abaI*, *abaR*) (Choi et al., 2009; Lannan et al., 2016; Ahmad et al., 2023). However, slight variations in VF combinations were observed between these two Malaysian community studies. Notably, the biofilm gene combination we termed BIO-Profile 1 (**Figure 1**) was dominant in our isolates but absent in those from the Segamat community (Muzahid et al., 2023). Despite these differences, the presence of numerous shared VFs underscores their potential role in supporting the persistence of the *A. baumannii* isolates in their niche and their capacity to cause infection.

**Figure 1:**
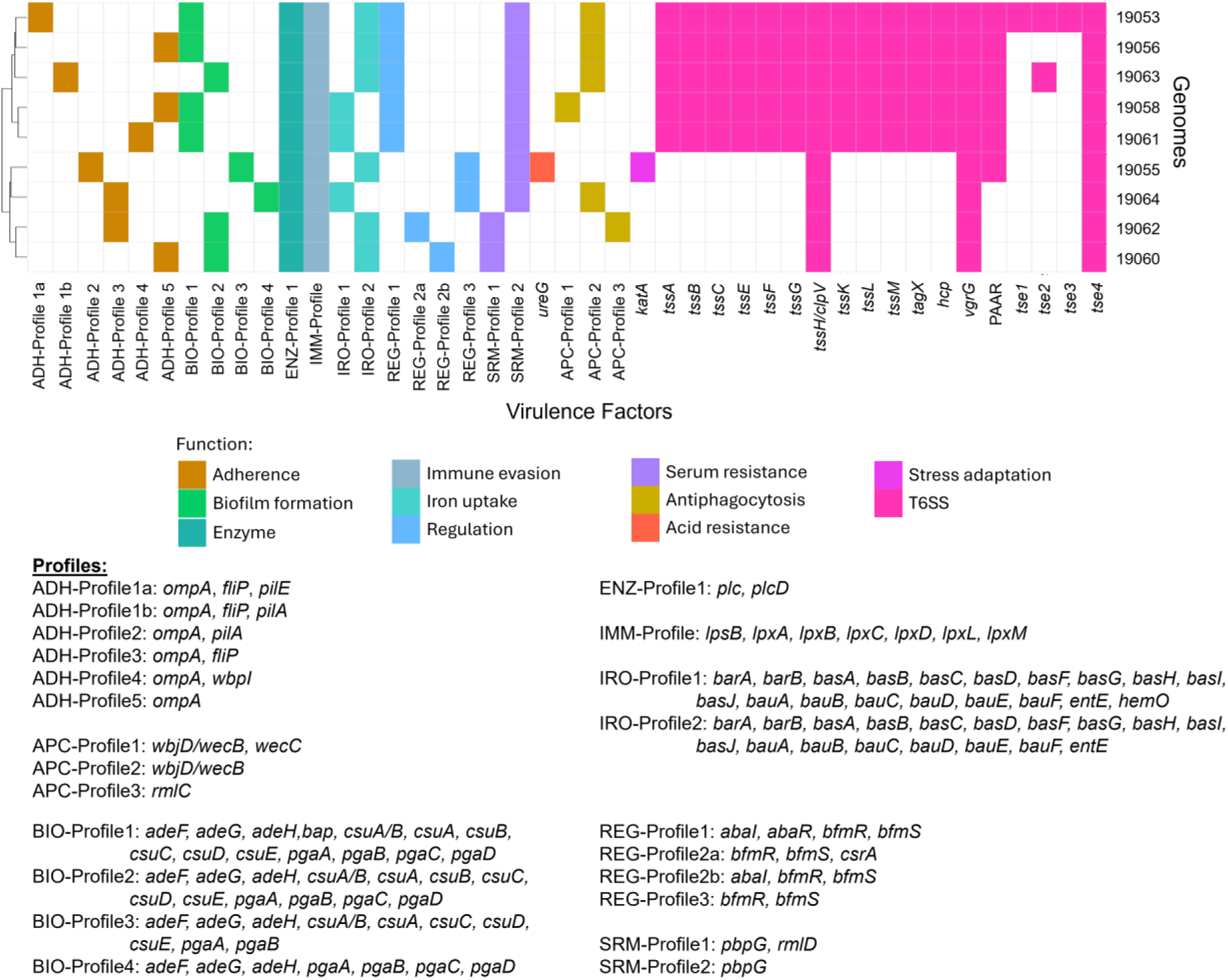
Profiles of various virulence factors (VFs) identified from the Orang Asli *A. baumannii* genomes (*n* = 9). The profiles were abbreviated according to their biological functions: adhesion (ADH), anti-phagocytosis (APC), biofilm formation (BIO), phospholipase enzyme (ENZ), immune evasion (IMM), iron uptake (IRO), regulation (REG) and serum resistance (SRM); singleton genes were labelled as is.

Survival of *A. baumannii* in harsh environmental conditions requires mechanisms for the acquisition of micronutrients, such as iron scavenging through the production of acinetobactin in iron-limiting environments (Lannan et al., 2016; Harding et al., 2018). The presence of iron uptake genes is thus a signature VF in *A. baumannii* and this was also observed in community isolates in this study and others (**Figure 1**) (Meumann et al., 2019; Muzahid et al., 2023). The combination of iron acquisition genes designated IRO-Profile2 (**Figure 1**) was also observed in the Segamat community isolates along with isolates from the Australian Northern Territory community (Meumann et al., 2019; Muzahid et al., 2023).

All nine *A. baumannii* isolates displayed the full complement of the *lps-lpx* genes (designated IMM-Profile; **Figure 1**), which are tagged as virulence factors that function in immune evasion. Deficiency in *lpxC* has been shown to cause the loss of the LPS layer in *A. baumannii* leading to the development of colistin resistance (Kamoshida et al., 2020). The full suite of the *lps-lpx* genes were found in the Segamat *A. baumannii* isolates, and this included hospital isolates that were identified as resistant to colistin (*n* = 2) and polymyxin B (*n* = 4); nevertheless, a more detailed analysis of possible mechanisms for polymyxin resistance, including mutations in the *lps-lpx* genes, was not presented (Muzahid et al., 2023).

The Type-6 Secretion System (T6SS) is utilized by *A. baumannii* to release toxic effector proteins into the neighbouring environment, offering a competitive advantage to the pathogen in multispecies environments (Carruthers et al., 2013) and also allowing *A. baumannii* to spread, invade and resist host immune responses (Shadan et al., 2023). The T6SS main cluster (T6MC), which encompasses the *tssA*, *tssB*, *tssC*, *tssD*/*hcp*, *tssE*, *tssF*, *tssG*, *tssH*/*clpV*, *tssK*, *tssL*, *tagX*, *vgrG* and PAAR genes (Fitzsimons et al., 2018; Lewis et al., 2019), was identified in five of the nine Orang Asli *A. baumannii* isolates (**Figure 1**). These genes are responsible for T6SS apparatus assembly, whereby TssA functions as the priming protein (also known as the cap), TssBC forms the sheath, TssD/Hcp the secretion tube with VgrG and PAAR proteins as the spike. The spike (i.e., VgrG and PAAR) teams with the wedge (i.e., TssK and TssEFG) to form the baseplate (Fitzsimons et al., 2018; Marazzato et al., 2022). The structure was then supported by the membrane complex formed by TssJ, TssM and TssL proteins connecting between inner and outer membrane (Fitzsimons et al., 2018; Marazzato et al., 2022). The T6SS found in the five *A. baumannii* genomes (**Figure 1**) was further identified as T6SS-1A (i.e., 19053, 19056 and 19063) and T6SS-1B (i.e., 19058 and 19061), according to the classification of (Repizo et al., 2019). These five *A. baumannii* isolates also encode the Tse4 effector (**Figure 1**), which function as an amidase (Lewis et al., 2019; Repizo et al., 2019). Other effectors were also present in the Orang Asli isolates, with *A. baumannii* 19063 and 19053 encoding an additional Tse2 (predicted to function as a DNase), while *A. baumannii* 19053 also encodes additional Tse1 (predicted lipase producer), and Tse3 (effector of unknown function) (Lewis et al., 2019; Repizo et al., 2019). In comparison, only one of the Segamat community isolate, *A. baumannii* C-98, harboured the complete T6SS whereas the hospital isolates contained the full suite of T6SS genes (Muzahid et al., 2023). Majority of the *A. baumannii* hospital isolates from Terengganu also harboured the full T6MC (*n* = 94/126; or 74.6%) (Din et al., 2025). It is thus tempting to speculate that the Orang Asli *A. baumannii* isolates were better able to survive in a multispecies environment with also the capacity to invade and colonise their host, should the opportunity arise; however, such possibilities would require further experimental investigations.

### 3.5 The Orang Asli A. baumannii isolates were genetically diverse

Community isolates of *A. baumanniii* represents a pool of unexplored genomes when compared to the more well-studied clinical isolates. Therefore, many *A. baumannii* isolates presented in community studies belonged to novel STs that have yet to be classified into any clonal complexes (CCs). We pooled together the *A. baumannii* genomes obtained from the Orang Asli in this study along with the genomes obtained from faecal samples of the community in the town of Segamat (Muzahid et al., 2023) for pangenome analysis. The analysis showed genetic diversity among these community isolates from the 5,277 cloud genes identified (53.84%) when compared to the 2,273 core genes (23.19%) (Figure 2). The lower ratio of core genes indicated low homogeneity between the *A. baumannii* community isolates, highlighting the uniqueness of the bacterium in each population. Even within the Segamat population itself, the higher diversity of the community *A. baumannii* isolates was apparent, as compared to the hospital isolates from the same town (Muzahid et al., 2023). Of interest, there did not seem to be any apparent geographical clustering between the two populations (**Figure 2**), and this was evident when examining the core genome phylogenetic tree that was generated using both community and clinical isolates of *A. baumannii* from Malaysia (**Figure 3**). Segamat is a township in the state of Johor and is approximately 390 km to the south of the state of Terengganu where the sampling was carried out in the Orang Asli rural settlements.

**Figure 2:**
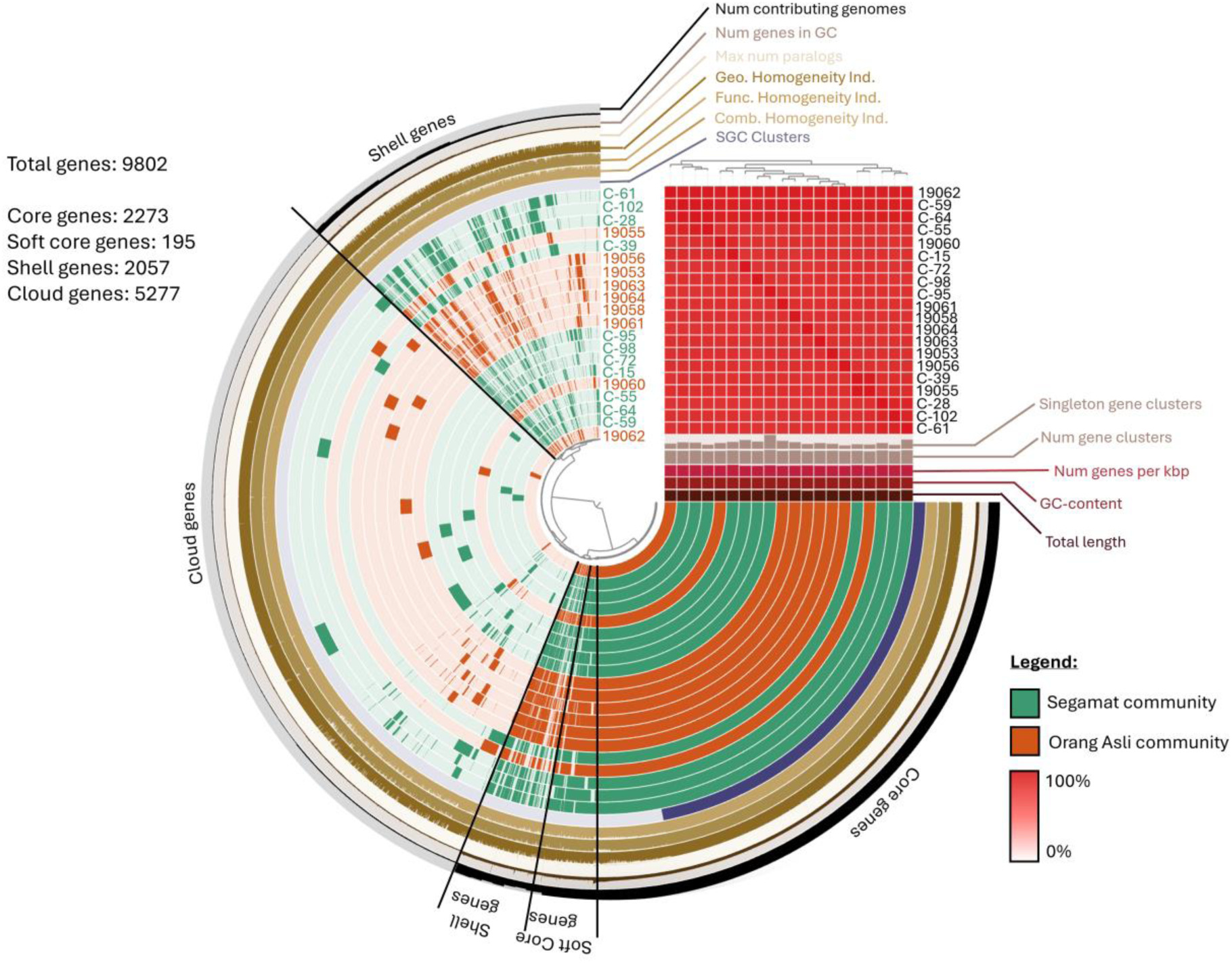
Pangenome analysis of community *A. baumannii* isolates from Malaysia that are currently available in the databases (*n* = 20). The orange-coloured tracks represent the Orang Asli community *A. baumannii* from this study (*n* = 9), whereas the green-coloured tracks represent the Segamat community strains (*n* = 11; isolates with the prefix “C”) that were previously published (Muzahid et al., 2023). Heatmap on the top right corner presents the average nucleotide identity (ANI) of *A. baumannii* from both communities, with all of them having >97% identity.

**Figure 3:**
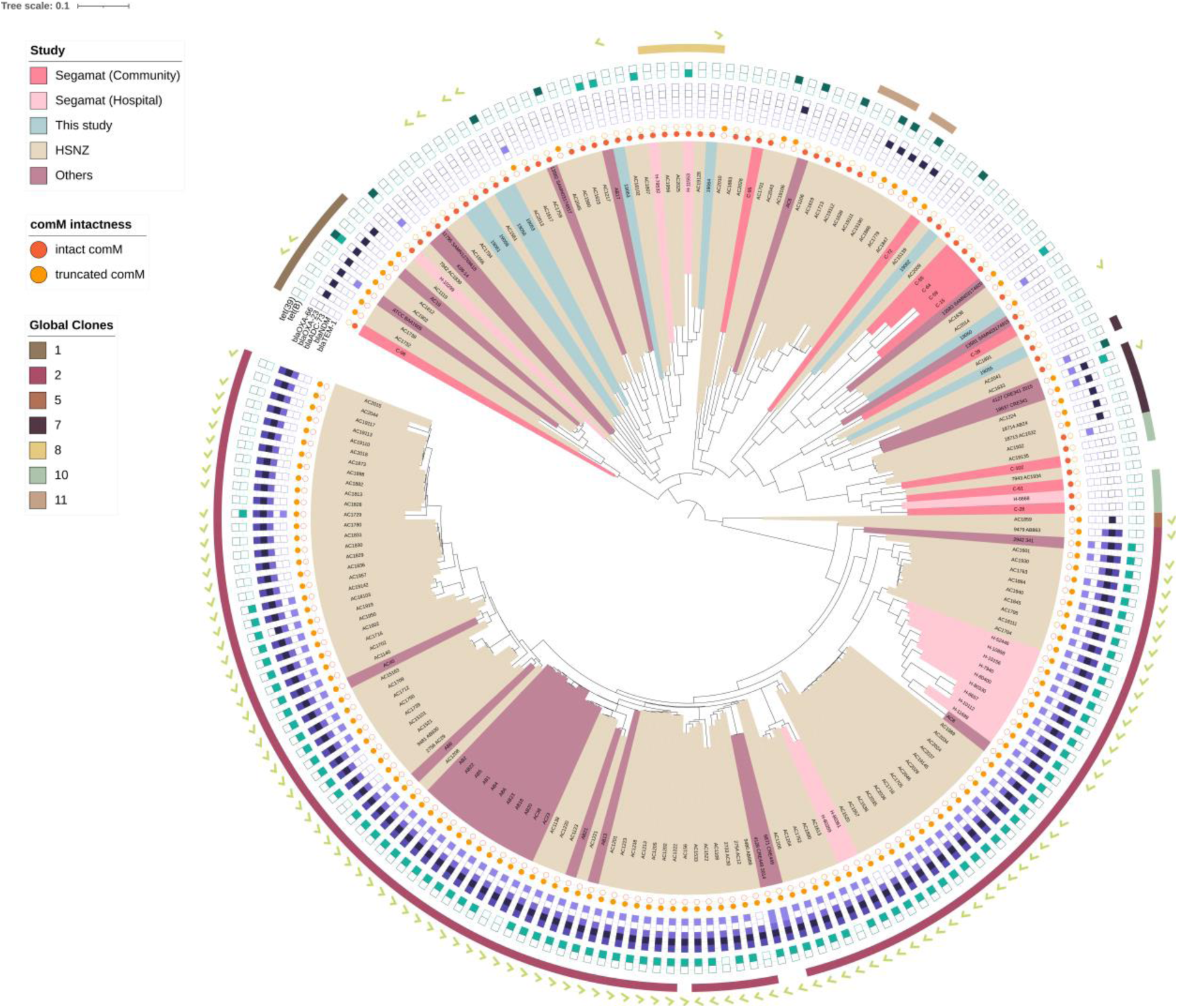
Midpoint-rooted maximum likelihood phylogenetic tree of all Malaysian *A. baumannii* genomes that are published [i.e., the current Orang Asli isolates, the Segamat hospital and community isolates described by (Muzahid et al., 2023), and the ten-year HSNZ isolates reported by (Din et al., 2025)] along with *A. baumannii* genomes originating from Malaysia that are found in PubMLST (as of 30 January 2025). Their categorization into the various Global Clone (GC) lineages are indicated. The presence of predominant carbapenemase genes from each Ambler class – *bla*_TEM-1_ (Class A), *bla*_NDM_ (Class B), *bla*_ADC-73_ (Class C) and *bla*_OXA-66_ (Class D) – are shown in filled purple boxes while their absences are indicated as empty purple boxes. Similarly, the presence/absence of the tetracycline resistance genes *tet(B)* and *tet(39)* are indicated in green boxes. Interruption of the *comM* gene, which is the hotspot for the insertion of the AbaR-type resistance islands, is shown as filled orange and light orange circles (labelled as intact *comM* and truncated *comM*, respectively). Light green tick marks at the outermost circle represent the presence of plasmids of the Rep_3 family (R3-type).

A maximum-likelihood phylogenetic tree was generated from the core genome alignment of 199 Malaysian *A. baumannii* genomes (**Figure 3**) and these included the genomes from this study, the Segamat study (both community and hospital isolates) (Muzahid et al., 2023), 126 genomes from a ten-year collection of isolates from Hospital Sultanah Nur Zahirah (HSNZ), the main tertiary hospital in Terengganu (Din et al., 2025), and other clinical isolates of *A. baumannii* obtained from the PubMLST database. Hospital isolates of *A. baumannii* were predominantly ST2_Pas_ which were categorized under the Global Clone 2 (GC2) lineage (Din et al., 2025) and this is clearly evident in the phylogenetic tree where they are clustered in a distinct clade (**Figure 3**). GC2 is the predominant *A. baumannii* lineage globally (Nigro and Hall, 2018) but the basis of this predominance is the overwhelming majority of sequenced isolates were from the hospitals (Shelenkov et al., 2023). As mentioned earlier, the Orang Asli *A. baumannii* genomes were scattered throughout the phylogenetic tree, as were the Segamat community isolates whereas most of the Segamat hospital isolates were clustered together in the GC2 clade, much like the HSNZ isolates. Interestingly, one Orang Asli isolate, *A. baumannii* 19064 (ST10_Pas_; ST585_Oxf_), was found to be a member of the GC8 lineage. Similarly, Muzahid et al. (2023) had reported that one of their community isolates from Segamat was identified as a member of the GC1 lineage. Apart from these exceptional cases, none of the community *A. baumannii* isolates belonged to any GC clusters. Additionally, the Orang Asli *A. baumannii* was also distinct from the Segamat community isolates with only *A. baumannii* 19062 distantly grouped with C-72 (**Figure 3**). Although majority of the Orang Asli isolates did not belong to any of the major Global Clones, we observed that they were interleaved with a few non-GC clinical isolates from HSNZ in neighbouring branches (**Figure 3**). It is possible that these community *A. baumannii* isolates were able to cause infections whenever the opportunity arises and thus, we see the relatively close genetic relationship between some of the Orang Asli isolates and the non-GC hospital isolates.

Apart from the distinct clustering of various GCs (Figure 3), the Malaysian clinical *A. baumannii* isolates also presented a different catalogue of dominant *bla* genes, which varies from the community *A. baumannii* resistome described earlier. The presence of *bla*_TEM-1_ (Ambler Class A), *bla*_NDM-1_ (Ambler Class B), *bla*_ADC-73_ (*bla*_ADC-1-like_; Ambler Class C), *bla*_OXA-23_ and *bla*_OXA-66_ (Ambler Class D) were recorded from almost all the Malaysian GC2 genomes, and likewise for the tetracycline resistance gene *tet(B)* (**Figure 3**). One distinctive feature of *A. baumannii* hospital isolates is the presence of large AbaR resistance islands that were often inserted within the chromosomal *comM* gene (Meumann et al., 2019). The intactness of the *comM* gene was investigated for all the Malaysian *A. baumannii* genomes presented here. Not surprisingly, the *comM* gene was interrupted in all the GC2 isolates whereas the proportion was 87.5% for GC1, 71.4% for GC7, and 66.7% for GC11 isolates (**Figure 3**). The non-GC *A. baumannii* genomes have a much lower percentage of interrupted *comM*, and within the nine Orang Asli isolates, only *A. baumannii* 19060 was identified with disruption of the *comM* gene (**Figure 3**). Nevertheless, the genes that were inserted within *comM* in *A. baumannii* 19060 were not associated with antimicrobial resistance but rather those associated with metabolism, regulatory genes and hypothetical proteins, much like what was described for the community-onset isolates from the Australian Northern Territory (Meumann et al., 2019).

### 3.6 Insertion sequence (IS) elements

A total of 17 ISs belonging to five families, namely IS*3*, IS*5*, IS*91*, IS*630* and IS*L3*, were identified from the Orang Asli *A. baumannii* genomes (**Supplementary Table 1**). IS*Aba43* of the IS*L3* family (Cameranesi et al., 2020) was found in all nine *A. baumannii* genomes, with at least one copy of the IS in each genome (**Supplementary Table 1**). Other IS families were found in fewer numbers with IS*Aba40*, IS*Aba57*, and IS*Aba63* of the IS*3* family found in one genome each, and the remaining IS elements identified found only in *A. baumannii* 19062 (**Supplementary Table 1**). There were few reports for most of these IS elements except IS*1006* which was identified in *A. baumannii* 19062 and is well-known for its association with plasmid-borne antimicrobial resistance regions (Harmer and Hall, 2021; Varani et al., 2021; Hall, 2022). Nevertheless, the copy of IS*1006* in *A. baumanii* 19062 was not found to be associated with any resistance genes.

### 3.7 Carriage of plasmids in A. baumannii from the Orang Asli community

Plasmids play an important role in the evolution of *A. baumannii*, being a major source for the dissemination of antibiotic resistance genes (Lam and Hamidian, 2024; Tobin et al., 2025). In a comprehensive survey of 439 mostly complete *A. baumannii* genomes, (Lam and Hamidian, 2024) reported that more than half (52%) contained one plasmid, 27% harboured two plasmids while seven genomes contained six to eleven plasmids. Plasmids of the Rep_3 family were by far, the most predominant and more than half of these plasmids carried antibiotic resistance genes (Lam et al., 2023; Lam and Hamidian, 2024; Tobin et al., 2025). Among the nine Orang Asli *A. baumannii* genomes, only one isolate, *A. baumannii* 19055, was without any detectable plasmids; three isolates (i.e., 19053, 19062 and 19064) harboured two plasmids each, while the remaining five isolates contained a plasmid each (**Supplementary Table 1**). Eight of these plasmids were small plasmids (i.e., <10 kb; (Lean and Yeo, 2017)), ranging in size from 2,178 bp to 8,837 bp. Out of these eight small plasmids, only one plasmid, p19053a, belonged to the Rep_1 family, specifically the R1-T6 type. p19053a was only 2,178 bp and harboured the *rep* gene and two other hypothetical open reading frames (ORFs) (**Supplementary Figure 1**). Lam and Hamidian (2024) (Lam and Hamidian, 2024) noted that the R1-type plasmids are typically 2 – 3 kb in size and encode only the replication initition protein along with one or two hypothetical proteins. None of these plasmids harboured AMR genes and p19053a follows the characteristics of the prototypical R1-type plasmid. The remaining seven small plasmids were of the Rep_3 family, specifically the R3-T5 (*n* = 4), R3-T13 (*n* = 2) and R3-T64 (*n* = 1) types

(**Supplementary Figure 1; Supplementary Table 1**). Two of these R3 types (i.e., R3-T5 and R3-T13) were reportedly among the most abundant R3 types globally, but their distribution were mainly linked to minor STs (Lam and Hamidian, 2024). Plasmid p19064a from the R3-T5 type was found in an ST10_Pas_ host (*A. baumannii* 19064), which was the only isolate from this study that belonged to the one of the known Global Clone lineages, GC8, but did not harbour antibiotic resistance genes. Among these small plasmids, only p19053b, which was also of the R3-T5 type, harboured the *tet(39)* tetracycline resistance gene (**Figure 4**). In contrast, *A. baumannii* clinical isolates from Malaysia, particularly those of the GC2 lineage, prevalently harboured an 8,731 bp plasmid we initially designated pAC12a (Lean et al., 2014; Lean and Yeo, 2017; Din et al., 2025) and which is identical to the pA1-1 plasmid (accession no. CP010782) that is harboured in a GC1 isolate obtained in 1982. This plasmid was previously typed as GR2 in an older scheme (Bertini et al., 2010) but has since been typed as R3-T1, and almost half of the members of this group were identical or nearly identical to pA1-1 (Lam et al., 2023). This attested to the long association of this plasmid with the GC1 and GC2 lineages but isolates of other STs have also been found that harbour this plasmid albeit at a much lower frequency (Lam et al., 2023; Din et al., 2025). A characteristic feature of this plasmid is the presence of mobile p*dif* modules containing the *sel1*, *abkBA* toxin-antitoxin genes, and a gene encoding a TonB-dependent receptor (Lean and Yeo, 2017; Lam et al., 2023). Notably, this plasmid type was absent from all *A. baumannii* isolates obtained from the Orang Asli population and the Segamat community, suggesting a likely specific association with clinical strains.

**Figure 4:**
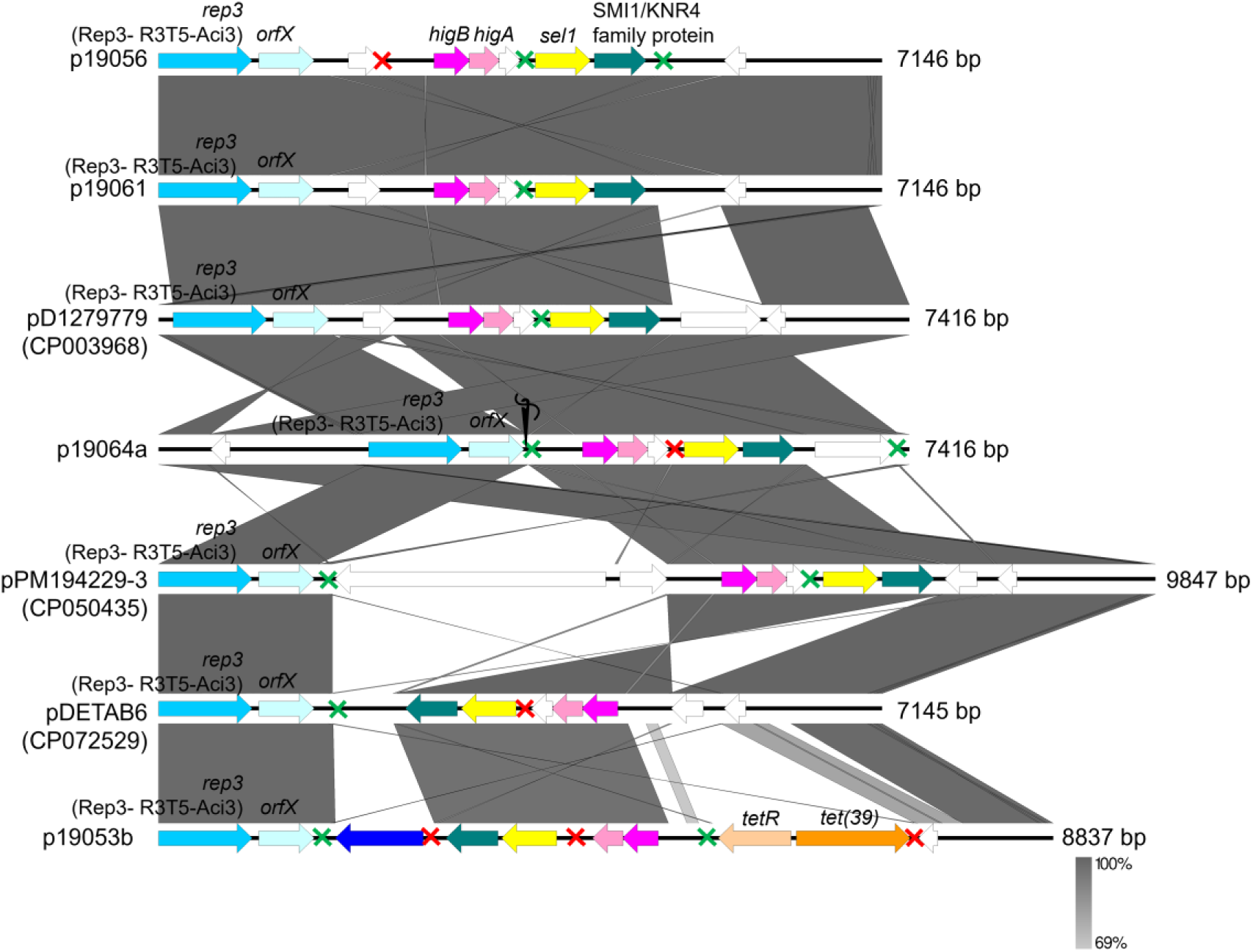
Comparisons of the R3-T5-type plasmids found in the Orang Asli *A. baumannii* isolates with similar R3-T5-type plasmids in the database curated by (Lam et al., 2023). The Rep_3 family replication initiation protein-encoding gene is indicated in blue arrows while its downstream gene designated *orfX* (Lam et al., 2023) is depicted in light blue arrows. The *higBA* toxin-antitoxin system is indicated in pink arrows (darker pink for the *higB* toxin and lighter pink for the *higA* antitoxin genes); the *tet(39)-tetR* tetracycline resistance gene pair is shown as orange arrows; the *sel1* gene encoding a Sel1-repeat protein is shown as yellow arrows; and the gene encoding the SMI1/KNR4 family protein is depicted as green arrows. White arrows indicate hypothetical open reading frames. The p*dif* sites are marked with green and red crosses representing XerC/D and XerD/C recognition sites, respectively. Needle and thread icon shown for the p19064a plasmid map indicated that the plasmid was a composite that was stitched together from two separate contigs.

Lam and Hamidian (2024) reported that almost half of the R3-type plasmids were not associated with AMR determinants. The genetic structures of the R3-T5 and R3-T13-type plasmids identified in this study (*n* = 6; **Figure 4** and **Supplementary Figure 1**) showed the absence of AMR genes from five of them, except for the *tet(39)-tetR* genes in p19053b. The *tet(39)-tetR* genes are located within a mobile p*dif* module usually found in *Acinetobacter* plasmids (Blackwell and Hall, 2017; Lean and Yeo, 2017). A p*dif* module typically comprises one or two related genes flanked by p*dif* sites, which resemble the chromosomal *dif* site involved in site-specific recombination. These 28 bp p*dif* sites contain binding regions for the recombinases XerC and XerD, similar to the chromosomal *dif* site near the bacterial chromosome terminus. The term p*dif* was used to differentiate these plasmid-associated sites from chromosomal *dif* sites (Blackwell and Hall, 2017; Castillo et al., 2017; Balalovski and Grainge, 2020). Three other p*dif* modules were uncovered from p19053b, and these carry a nucleotide-binding protein-encoding gene, *higBA* toxin-antitoxin genes, and a *sel1* gene along with a gene encoding the SMI1/KNR4-family protein (**Figure 4**). Interestingly, the gene arrangement of the *higBA*, and *sel1*-SMI1/KNR4-family p*dif* modules were observed in almost all the R3-T5-type plasmids identified in this study and also extends to the R3-T13-type plasmids (i.e., p19058 and p19063) (**Figures 4** and **5; Supplementary Figure 1**).

**Figure 5:**
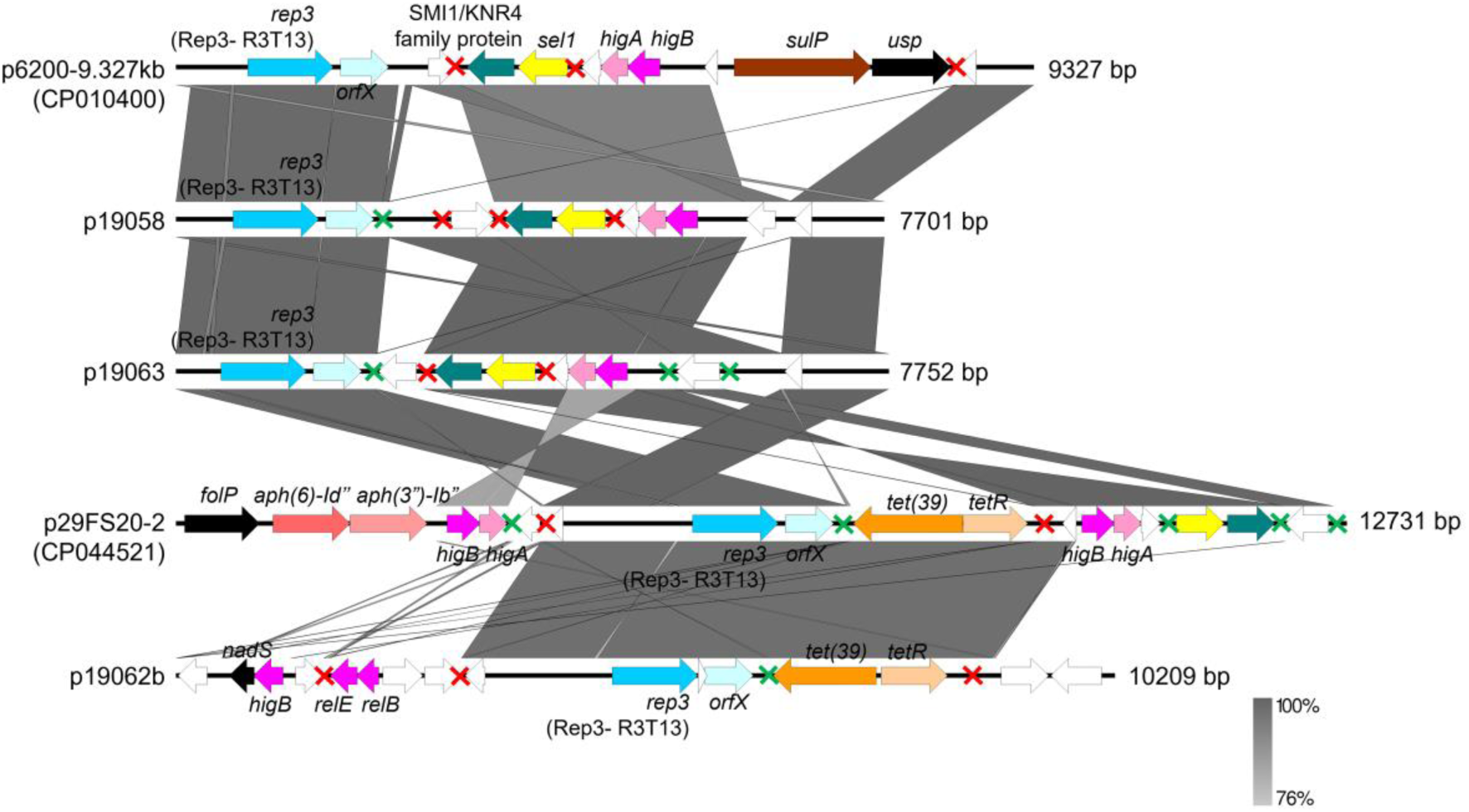
Comparative linear map of R3-T13-type plasmids identified from the Orang Asli *A. baumannii* with closely related R3-T13-type plasmids found in the plasmid database presented by (Lam et al., 2023). Coloured arrows representing genes and red-green crosses representing p*dif* sites were labelled as indicated in the previous Figure. The only antibiotic resistant R3-T13-type plasmid found in the database was p29FS20-2 which was harboured by a tigecycline-resistant *A. baumannii* isolated from faeces in Guangdong, China (accession no. CP044521.1). Plasmid p19062b shared the *rep3-orfX* and the *tet(39)-tetR* genes with p29FS20-2.

Two plasmids of the R3-T5-type, namely p19056 and p19061, were nearly identical, with both at 7,146 bp (**Figure 4**). They were closely related to the pD1279779 plasmid (accession no. CP003968.1) which was slightly larger at 7,416 bp with an additional hypothetical protein found downstream of the SMI1/KNR4-family gene (**Figure 4**). Plasmid p19064a also appeared to be closely related to pD1279779 but the p19064a sequence was actually stitched together from two separate contigs (**Figure 4**), and whether there are additional genes that are lost in between these two contigs is not known. Interestingly, the host for pD1279779 was *A. baumannii* D1279779, which was isolated from a community-acquired bacteraemia patient in Northern Australia and typed as ST267_Pas_/ST942_Oxf_ (Farrugia et al., 2013). Based on the collection of *Acinetobacter* plasmids curated by (Lam et al., 2023), the R3-T5-type plasmids identified in this study were related to pAba10324b, pPM194229-3 and pDETAB6, besides pD1279779. The p19053b plasmid, which harboured the *tet(39)-tetR* genes, is thus the sole R3-T5-type plasmid which encode AMR determinants. This plasmid is more closely related to pDETAB6, which harboured two hypothetical ORFs instead of the *tet(39)-tetR* genes in p19053b (**Figure 4**).

AMR genes are also a rarity among the R3-T13-type plasmids. A search in the *Acinetobacter* plasmid repository posted by (Lam et al., 2023) led to the discovery of p29FS20-2 (accession no. CP044521.1) as the sole carrier of *tet(39)-tetR*, and the aminoglycoside resistance genes *aph(3’’)-Ib* and *aph(6)-Id* in this plasmid type. Two of the three R3-T13-type plasmids identified in this study, p19058 and p19063, were not associated with AMR determinants; however, p19062b was the only R3-T13-type plasmid that harboured the *tet(39)-tetR* gene pair (**Figure 5**). Similar to p19053b, the *tet(39)-tetR* genes in p19062b was also located within a p*dif* module, a feature that was also observed in p29FS20-2 (**Figure 5**). Plasmids p19058 and p19063 were more closely related to another R3-T13-type plasmid, p6200-9.327kb, but this plasmid contained additional genes encoding sulphate permease (*sulP*) and a universal stress protein (*usp*) (**Figure 5**). One of the interesting observations of note regarding these plasmids is the almost universal presence of the *rep_3-orfX* backbone, the *higBA*, and *sel1-*SMI1/KRN4 family modular arrangement of genes, covering also the R3-T5-type plasmids (**Figures 4** and **5**). The sole exception to this is plasmid p19062b where the *sel1*-SMI1/KRN4 p*dif* module was absent, and the putative toxin-antitoxin module (identified as *relBE* by TADB 3.0) was distantly related to the *higBA* module usually found in the other similar plasmid-types (**Figure 5**).

The other plasmids identified in the Orang Asli *A. baumannii* genomes are p19062a of the R3-T64 type, p19060 of R3-T26, and p19064b of R3-T27 type (**Supplementary Figure 1**). These plasmid types are rarely encountered when compared to the R3-T5 and R3-T13 types (Lam et al., 2023). None of these plasmids harbour AMR determinants.

A major limitation in this study is that these *A. baumannii* genomes were sequenced using the Illumina short-read platform, which does not allow for complete genome assembly. Therefore, some of the plasmid architecture presented here should be taken with caution as we could not ascertain if there are genes or genetic elements that are lost in the assembly of the draft genomes. This limitation is particularly relevant for plasmid p19064a which was assembled across two separate contigs. Nevertheless, the fact that some of these plasmids have similar counterparts from clinical *A. baumannii* isolates is of concern as they could serve as vehicles for the dissemination of AMR genes. The discovery of the *tet(39)-tetR* gene pair within a mobile p*dif* module in two distinct plasmids, p19053b and p19062b, underlines this likelihood and highlights the importance of continued genomic surveillance particularly among the community.

## 4 Conclusions

*Acinetobacter baumannii* and its antimicrobial resistance mechanisms have long been central in the global fight against superbugs, as understanding these traits is essential for improving treatment strategies. Over the decades, numerous reports have highlighted the alarming rise in carbapenem-resistant, MDR and even PDR *A. baumannii* strains in clinical settings (Akeda, 2021; Chen et al., 2023), suggesting that that treatment practices themselves may contribute to the development of resistance (De Blasiis et al., 2024). In contrast, *A. baumannii* isolates from remote communities, such as the indigenous Orang Asli population studied here, exhibited lower levels of antibiotic resistance. However, data on community-derived isolates remain scarce when compared to clinical isolates (Lam and Hamidian, 2024).

Despite their relative antibiotic susceptibility, these community-associated *A. baumannii* isolates still harbour a range of virulence factors (VFs) and mobile genetic elements, including plasmids and insertion sequences (Meumann et al., 2019; Muzahid et al., 2023), features well-documented in hospital-derived strains. Notably, one isolate from this study belonged to the Global Clone 8 (GC8) clinical lineage, while (Muzahid et al., 2023) previously identified a Global Clone 1 (GC1) isolate in a more urbanised community setting in Malaysia. Moreover, the phylogenetic interleaving of these community isolates with certain non-GC hospital strains suggests that they possess the capacity to cause infections and acquire resistance traits, akin to their clinical counterparts. The presence of shared genomic features in *A. baumannii* from a presumptive antibiotic-naïve environment underscores the pathogen’s inherent capacity for persistence and colonization, even in healthy individuals. Of concern is the detection of two *A. baumannii* isolates from this study that harbours tetracycline resistance genes in mobile p*dif* modules located in distinct plasmids with similarities to those isolated from clinical strains. This finding suggests the potential for further acquisition of resistance determinants and serves as a cautionary signal against the unregulated introduction of antibiotics into vulnerable populations. Healthcare practitioners working with remote communities can use these insights to make more informed treatment decisions, potentially reducing the emergence and spread of AMR (Yau et al., 2021). Therefore, expanding genomic surveillance to include community-derived *A. baumannii* strains, even from remote indigenous tribes is indeed a useful endeavour. Although this is a road less taken, the knowledge obtained particularly tracking the shifts in known and novel STs, will be invaluable for understanding the pathogen’s broader epidemiological dynamics and informing future public health strategies.

## Data availability

The nine *Acinetobacter baumannii* genomes have been deposited in the National Center for Biotechnology Information (NCBI)’s Genomes database under BioProject no. PRJNA1258958.

## Acknowledgements

We would like to thank Prof Ramle Abdullah, Dr Hafis Simin and their Orang Asli research group for their kind assistance. Our gratitude also to Dr Salwani Ismail, Dr Nor Iza A. Rahman, Dr Mohd Sayuti Razali, Dr Nor Kamaruzaman Esa, Dr Salman Amiruddin, and Dr Chew Ching Hoong for their kind support and assistance during the sampling trips. We would also thank the Department of Orang Asli Affairs and Development (JAKOA) for providing access to the Orang Asli villages in Terengganu. Our gratitude goes to the communities who partook their support and participation in this study.

## Ethics approval and consent to participate

Ethical approval for isolates taken in Peninsular Malaysia was provided by Universiti Sultan Zainal Abidin (UniSZA) Ethics Committee: approval no. UniSZA/C/1/UHREC/628–1(85) dated 27 June 2016, the Department of Orang Asli Affairs and Development (JAKOA): approval no. JAKOA/PP.30.052Jld11[42], and by the University of Southampton Faculty of Medicine Ethics Committee (Submission ID: 20831). Written informed consent was taken with parents/guardians providing consent for those < 18 years old.

## Author contributions

C.C.Y., S.C.C., and D.W.C. conceived the study. S.C.C., D.W.C., and C.C.Y. secured funding. C.C.Y and A.G.A. were responsible for study planning and visits in Malaysia. D.E.M. and A.G.A. conducted the visits for sampling. R.A. and D.E.M. undertook microbiological culture and sample preparation for sequencing. S.S.L., A.G.A., D.W.C., and C.C.Y. undertook data analysis. S.S.L. and C.C.Y wrote the manuscript. All authors reviewed the manuscript and approved the submitted version.

## Consent for publication

All authors have provided consent for publication.

