## Supplementary Figure 1 for "The road less travelled: Exploring the genomic characteristics and antimicrobial resistance potential of *Acinetobacter baumannii* from the indigenous Orang Asli community in Peninsular Malaysia"

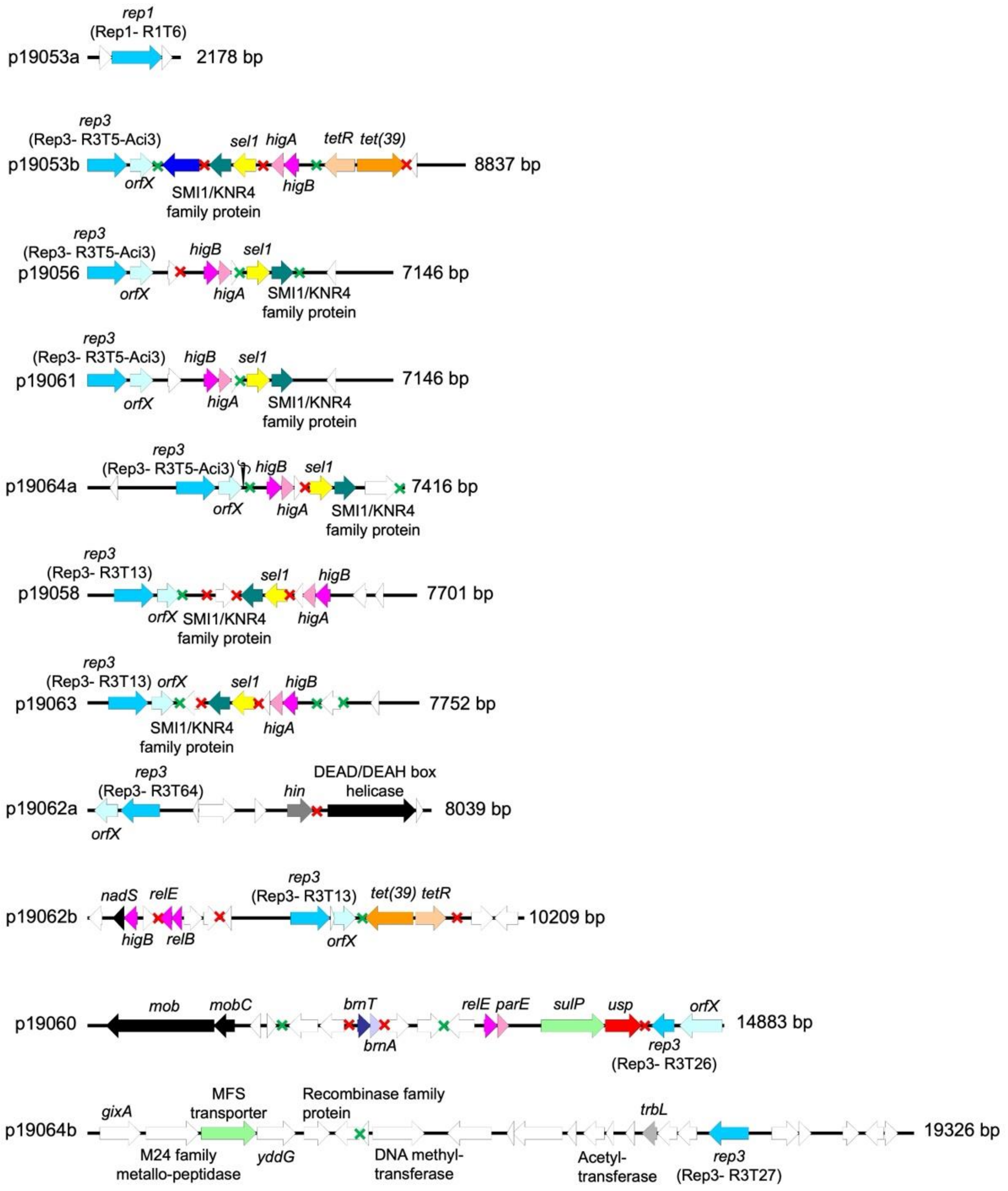


**Supplementary Figure S1**: Linear genetic maps of all plasmids identified in the Orang Asli community *A. baumannii* isolates. Majority of the plasmids were of the Rep_3 family with only plasmid, p19053a, belonging to the Rep_1 family. The *rep*-encoded replication initiation protein gene is depicted as blue-coloured arrows with its Rep type labelled while its downstream gene, recently designated *orfX* (Lam et al., 2023), is shown as light blue arrows. White coloured arrows represent genes encoding hypothetical proteins. Other coloured arrows are as in the legends to **Figures 4** and **5** in the main text. p*dif* sites are depicted as green (XerC/D) and red (XerD/C) crosses. Note that plasmid p19064a is a composite of two contigs that were stitched together as signified by the needle and thread icon above its linear map.
